# Integration of intra- and inter-sexual selection signaling

**DOI:** 10.1101/2020.05.14.088518

**Authors:** Courtney R. Garrison, Raphaël Royauté, Ned A. Dochtermann

## Abstract

Sexual selection can drive the evolution of dramatic morphological and behavioral signals. This selection acts on both specific components of signals and overall signals that combine multiple sources of information. By studying the structure and variability of signals and their components we can improve our understanding of how sexual selection operates. Signal integration can be understood through the lens of classical signaling hypotheses or more recently defined systems approaches. Using crickets (*Acheta domesticus*), we evaluated competing hypotheses about signal integration and how observed patterns of signal integration fit into both systems approaches and classic signaling hypotheses. We measured three call types of 127 male crickets multiple times for a total of 930 observations. We found evidence for an underlying integrated signaling syndrome from which both intra- and intersexual signals stemmed. This syndrome was also affected by mass, suggesting honest signaling in the species. The presence of an integrated syndrome demonstrates that intra- and intersexual signals are incorporated in a redundant signal strategy in *Acheta domesticus*. This support for honest and redundant signaling is also consistent with a systems framework description of signals as degenerate and functionally modular—demonstrating one way in which classic hypotheses can be integrated with modern systems approaches.

## Introduction

Sexual selection results from differing reproductive success due to among-individual variation in traits affecting reproductive success. Traits most typically shaped by sexual selection include male and female genital morphology and sexual signals such as color patterns, vocalizations, and courtship displays that relay information to the opposite sex (Andersson and Simmons 2006). Sexual signaling is important across taxa (e.g. Harrison et al. 2013, Moreno-Gomez et al. 2015) and sexual selection frequently leads to the evolution of increasing signal complexity over time (Buchanan et al. 2003, Spencer et al. 2003, Woodgate et al. 2012). Signals can also be produced for functionally distinct interactions, with many species producing signals specific to intra-or intersexual communication. Because sexual selection can act on specific components of signals and across functionally distinct signals (Hedrick 1986, Hedrick and Weber 1998, Buchanan et al. 2003, Spencer et al. 2003, Woodgate et al. 2012), studying signal structure and signal production can provide valuable insights regarding the action of sexual selection.

The relationship among signal components, i.e. the pattern of phenotypic integration (Pigliucci 2003), has important implications for how signals might be interpreted by receivers. This interpretation by receivers provides the framework for a number of classic hypotheses that have facilitated our general understanding of animal communication (Table 1). For example, the “redundant signal” hypothesis, states that signals convey redundant, shared information to the receiver (Johnstone 1996). As a modification of the redundant signal hypothesis, individuals might have a general signaling phenotype which causally influences the repertoire of signals produced. Such a general signaling phenotype can represent an honest signal, providing accurate information about condition to a receiver (Zahavi 1975, Schluter and Price 1993, Berglund et al. 1996). These non-mutually exclusive hypotheses necessarily require strong patterns of integration (i.e. strong correlations) among signal components. Alternatively, if signals and their components show weak patterns of integration, each signal may provide receivers with distinct information about the signaler, a form of the “multiple-message” hypothesis (Johnstone 1996). If signal components are uncorrelated, this can be interpreted as support for the multiple message hypothesis since the uncorrelated signals are providing independent information about the signaler.

**Table 1.** Classic signaling hypotheses and corresponding systems approach labels, descriptions, examples, and suggested integrated terminology

| **Classic Signaling Hypothesis** | **Systems Approach Hypothesis** | **Description** | **Example (in *Acheta domesticus*)** |
| --- | --- | --- | --- |
| Redundant Signals | Degeneracy | Signal components serve a similar function | Call frequency and pulses per chirp provide the receiver information about male body size |
| Redundant Signals | Functional Modularity | Signal components serve a similar function, while overarching call types/modules serve similar functions | Call frequency and pulses per chirp provide the receiver information about male body size, advertisement and aggression calls both communicate information about signaler's size |
| Multiple Messages | Functional Modularity | Signal components serve a similar function within a call type/module, while the overarching call types serve different functions | Call frequency and pulses per chirp provide the receiver information about male body size, but aggression and courtship calls are used for different functions |
| Honest Signaling | Degeneracy | Signal components serve a similar function and are causally affected by condition or "quality" | Call frequency and pulses per chirp provide the receiver information about male body size, with mass affecting both signal components |
| Honest Signaling & Redundant Signals | Degeneracy & Functional Modularity | Signal components serve a similar function, while overarching call types/modules serve different functions, all of this is causally affected by condition or "quality" | Call frequency and pulses per chirp provide the receiver information about male body size, with mass affecting both signal components, while aggression is used towards other males and courtship is used towards females |

Both the redundant signaling and multiple message hypotheses have been supported in a variety of taxa. For example, male eland antelopes (*Tragealphus oryx*) exhibit redundant signaling via facemask darkness, frontal brush size and body greyness—traits correlated with androgen levels and aggression (Bro-Jørgensen et al. 2008). Meanwhile, support for the multiple message hypothesis has also been frequently observed. As one example, bowerbird males (*Ptilonorhynchus violaceus*) signal health and condition with both bower characteristics and plumage coloration (Doucet and Montgomerie 2003). Importantly, the redundant signaling and multiple message hypotheses are not entirely mutually exclusive: while facemask darkness, brush size and greyness indicates aggression in the aforementioned male eland antelopes, independent signals—such as a characteristic knee-click—provides independent information related to fighting ability (Bro-Jørgensen et al. 2008).

Recently, Hebets et al. (2016) proposed a systems approach to understanding communication. This framework stresses that an entire signaling system should be evaluated simultaneously (Hebets et al. 2016). The power of this proposal vis-à-vis sexual selection is that it properly recognizes the entire signal repertoire of an organism, and all the components of that repertoire, as simultaneously affecting reproductive success and therefore fitness. This systems approach formalizes the description of how signals interact both for a single mode of communication where multiple signals are employed or within a multimodal communication framework (sensu Hebets and Papaj 2005). According to this systems approach, signaling can be categorized to one of four different system designs (Table 1; Hebets et al. 2016, Rosenthal et al. 2018): 1) Redundancy, where structure and function are shared among signal components, this is shown by repetition of a song or display. 2) Degeneracy, where different signals or signal components serve similar functions. 3) Pluripotentiality, one signal or component of a signal serves multiple functions in one display (e.g. both intra and intersexual functions) and 4) Modularity, when subsets of signals form linked structural or functional clusters (Hebets et al. 2016, Rosenthal et al. 2018). With this framework Hebets et al. (2016) have provided animal signaling research an overarching approach to describing the relationship among signals and among the components of these signals.

Unfortunately, the classic hypotheses and the systems approach possess overlapping terminology—a signaling issue in its own right. For example, the classic redundant signals hypothesis which states that signals convey redundant, shared information to receivers (Johnstone 1996) is not the same as Hebets et al’s (2016) redundancy systems approach (i.e. the same signal repeated). While this overlap necessitates clarity in operational definitions, a more general issue is that while the systems approach clearly describes the structure of signals, the connection of the systems approach categorizations to sexual selection theory is less clear. One approach to resolve this issue is to recognize the connections between classical signaling hypotheses and the Hebets et al. (2016) framework to interpret the system structures (Table 1).

Here, we sought to determine how intra- and intersexual signals are integrated and used a hypothesis comparison approach to evaluate classic signaling hypotheses in the domestic cricket (*Acheta domesticus*). We simultaneously evaluated how the relationship among signal components fit within the systems approach of Hebets et al. (2016), allowing an integration of classic hypotheses with relevant sexual selection theory. Crickets are ideal for testing questions about signaling, signal complexity, and signal integration because they use the same physical structures in both intra- and intersexual signaling, exposing them to varying selection pressures based on receiver sex. Specifically, male crickets produce three distinct signals: an advertisement call used to attract females from long distances, a courtship call used to induce copulation by females after they have closely approached a male, and an aggression call used between males during agonistic encounters (Gray and Eckhardt 2001, Zuk et al. 2008). This combination of signals and their differing functions and targeted receivers allowed us to use the systems approach structure to describe if and how advertising, courtship, and aggression call types were integrated and which classic signaling hypothesis was supported by observed patterns of integration.

## Materials and Methods

### Diet and Rearing

Male nymph *A. domesticus* (obtained from Fluker’s Cricket Farm) were reared in plastic containers (34.6 x 21 x 12.4 cm). Each container housed around 10 nymphal crickets and was maintained at 32⍰ C on a 12:12 hr light cycle. Crickets were provided with egg carton pieces for housing and food and water *ad libitum* (Royauté and Dochtermann 2017, Royauté et al. 2019). Crickets were reared on one of four experimental diet regimes as part of a larger experiment (Royaut é et al. 2019), but these diets had no detectable effect on call components (Garrison 2017) or call covariances (Mantel tests’ r > 0.50, Table S1). Once crickets reached maturity, they were moved into individual containers (0.71-L) and fed an assigned diet and water *ad libitum*. Mature crickets were kept at a 12:12 hr light cycle at 25⍰ C

### Call Recording

*We* measured advertisement, courtship, and aggression calls repeatedly for a total of 127 male crickets, with a total of 930 calling trials (Table S2). We attempted to obtain three repeated measures per individual per call type but, due to natural mortality, some call types were recorded more frequently than others (Table S2). We used a repeated measures framework as the call components in crickets are influenced by both genetic and environmental effects (Hedrick 1988). By repeatedly measuring each call, we were able to estimate the among-individual variability of calls and their underlying components (Dingemanse and Dochtermann 2013) and among-individual correlations between call components (Dingemanse et al. 2012). These among-individual correlations estimate the combined contribution of genetic and long-term environmental effects on correlations, separate from temporary environmentally induced effects (Dingemanse and Dochtermann 2014.

To record advertisement calls, housing containers were surrounded by acoustic foam and USB audio recorders were placed on top of each individual container for 2 hours. Because females are not necessary to elicit advertisement calls in this species (Garrison 2017), males called over this period without a female cricket in the container.

To record courtship calls, a female cricket must be present with the male (Garrison, personal observation). Following Zuk et al. (2008), male crickets were introduced into a container the same size as those used for housing but containing only a USB audio recorder and a live female assigned at random. The females used in courtship trials were obtained from our laboratory stock collection so mating status of females was unknown but all males were virgins. Courtship call was then recorded for a period of 5-10 minutes. If a male failed to call within the first 5 minutes of the trial, the trial was stopped, and the male was removed and recorded as not calling. If a female attempted to copulate with (i.e. mounted) the focal male during the trial, recording was also stopped and mating was not allowed to be completed. All courtship calls were conducted at least 48 hours after the final advertisement calls had been recorded since potential contact with a female could alter the male’s long-range calling effort.

To record aggressive calls, focal males were placed in a novel container with a random male that had been muted by having its forewings removed. A pilot study showed that *A. domesticus* will produce aggressive calls towards other males without a female present as a stimulus (Garrison 2017), so females were not used to elicit aggressive calls. Aggressive calls were recorded for 5-10 minutes. Trials were ended when there was a clear winner (one cricket retreated). If a male failed to call within the first 5 minutes of the trial, the trial was stopped and the male was removed and recorded as not calling and tested again at a later date.

### Call Analysis

Each call was analyzed using the sound analysis programs Audacity and Avisoft. We measured 7 calling components: chirp rate, chirp duration, mean number of pulses per chirp, peak frequency, call amplitude, pulse rate and total time calling for all three call types (i.e. 21 total call components).

For advertisement calls, we analyzed the middle forty-five minutes of each recording. Courtship calls were more difficult to analyze as males intermittently produced advertisement calls during courtship. To properly analyze courtship calls, only sections of the courtship trial recordings that were exclusively courtship were analyzed. There is a visible difference in call structure between courtship and advertisement calls (Figure 1). A similar issue was encountered with aggression call, but aggression call is easily distinguished based on call waveforms (Figure 1).

**Figure 1.**
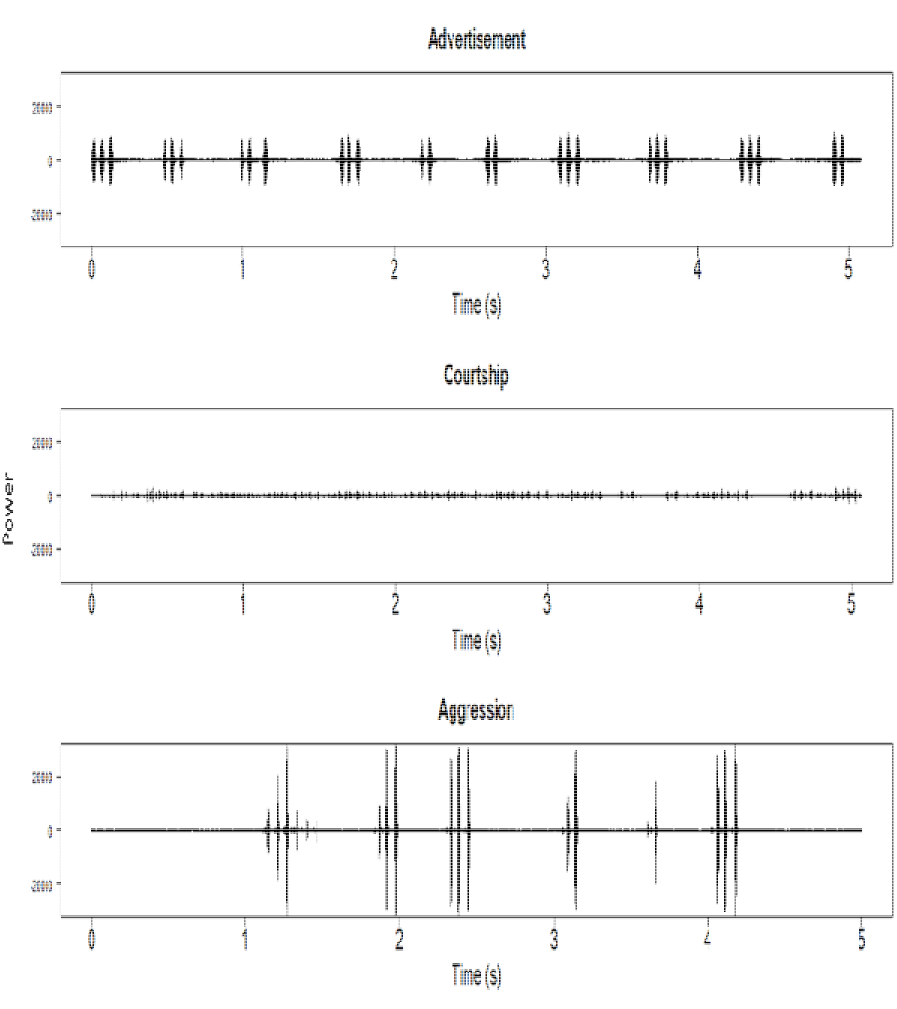
Waveform structures of five seconds of advertisement, courtship and aggression song from the same cricket. The y-axis represents the amplitude of the call.

## Statistical Analysis

### Repeatability by Call Type

As an initial exploratory analysis we estimated adjusted and unadjusted repeatabilities and the variation due to all random and fixed effects for each of the 21 call component using the `rptR` package (Nakagawa and Schielzeth 2010, Stoffel et al. 2017) in the R statistical language. In addition, which conspecific was present (male opponent for aggression call, female for courtship call), the chamber in which recording occurred (advertisement call), and the developmental box in which an individual was reared (all call types) were included as random effects and corresponding variances components estimated (Table S3).

### Comparing Hypotheses of Signal Integration

To test competing hypotheses of phenotypic integration we employed a two-step approach.

First, we estimated among- and within-individual covariance and correlation matrices (Dingemanse et al. 2012) for mass and the 9 call components that had previously been found to exhibit the highest repeatabilities in the previous exploratory univariate analyses (Table 2, Table S3). We chose the components with the highest repeatabilities as these will be least affected by temporary environmental effects (Falconer 1996). The among- and within-individual covariance and correlation matrices were estimated using multi-response mixed effects models with temperature (centered), repetition number, batch, and time of recording as well as age since maturation and diet type included as explanatory variables. Male identity was included as a random factor. The 10 response variables were mean and variance standardized to facilitate model fitting. The multi-response mixed effect model was then fit using the `MCMCglmm` package (Hadfield 2010) in the R statistical language. The model was fit with an MCMC chain with 1.3 × 10^6^ iterations, a 300000 burn-in period, and a thinning interval of 1000 and a prior that was flat for correlations. This chain length led to low autocorrelations and high MCMC effective sample sizes.

**Table 2.** Repeatabilities from univariate models for the traits chosen for the multi-response mixed-effects model with 95% confidence intervals

| **Signaling Trait** | **$\tau^a$** |
| --- | --- |
| **Advert Frequency** | 0.58 (0.45, 0.7) |
| **Advert Pulses per Chirp** | 0.46 (0.29, 0.60) |
| **Advert Chirp Rate** | 0.42 (0.27, 0.59) |
| **Aggression Frequency** | 0.39 (0.23, 0.54) |
| **Aggression Pulse Rate** | 0.31 (0.16, 0.50) |
| **Aggression Amplitude** | 0.22 (0.09, 0.42) |
| **Courtship Pulses per Chirp** | 0.32 (0.13, 0.55) |
| **Courtship Chirp Duration** | 0.32 (0.13, 0.55) |
| **Courtship Chirp Rate** | 0.29 (0.1, 0.53) |

<sup>a</sup>adjusted repeatability, calculated as $V_i/(V_i + V_w)$

Second, we specified structural equation models (SEMs) corresponding to nine hypothesized patterns of trait integration (Figure 2) prior to fitting the multi-response mixed effect model. We then fitted each of these SEMs to the posterior distribution of estimated among-individual correlation matrices (the MCMC analyses produced 1000 estimates of the correlation matrix) and evaluated SEMs in competition with each other based on Akaike Information Criteria values (AIC) following Araya-Ajoy & Dingemanse (2014). This combination of SEMs and AIC based model comparison approaches allows the testing of specific hypothesis of trait integration (Dochtermann and Jenkins 2007, Dingemanse et al. 2010, Araya-Ajoy and Dingemanse 2014).

**Figure 2.**
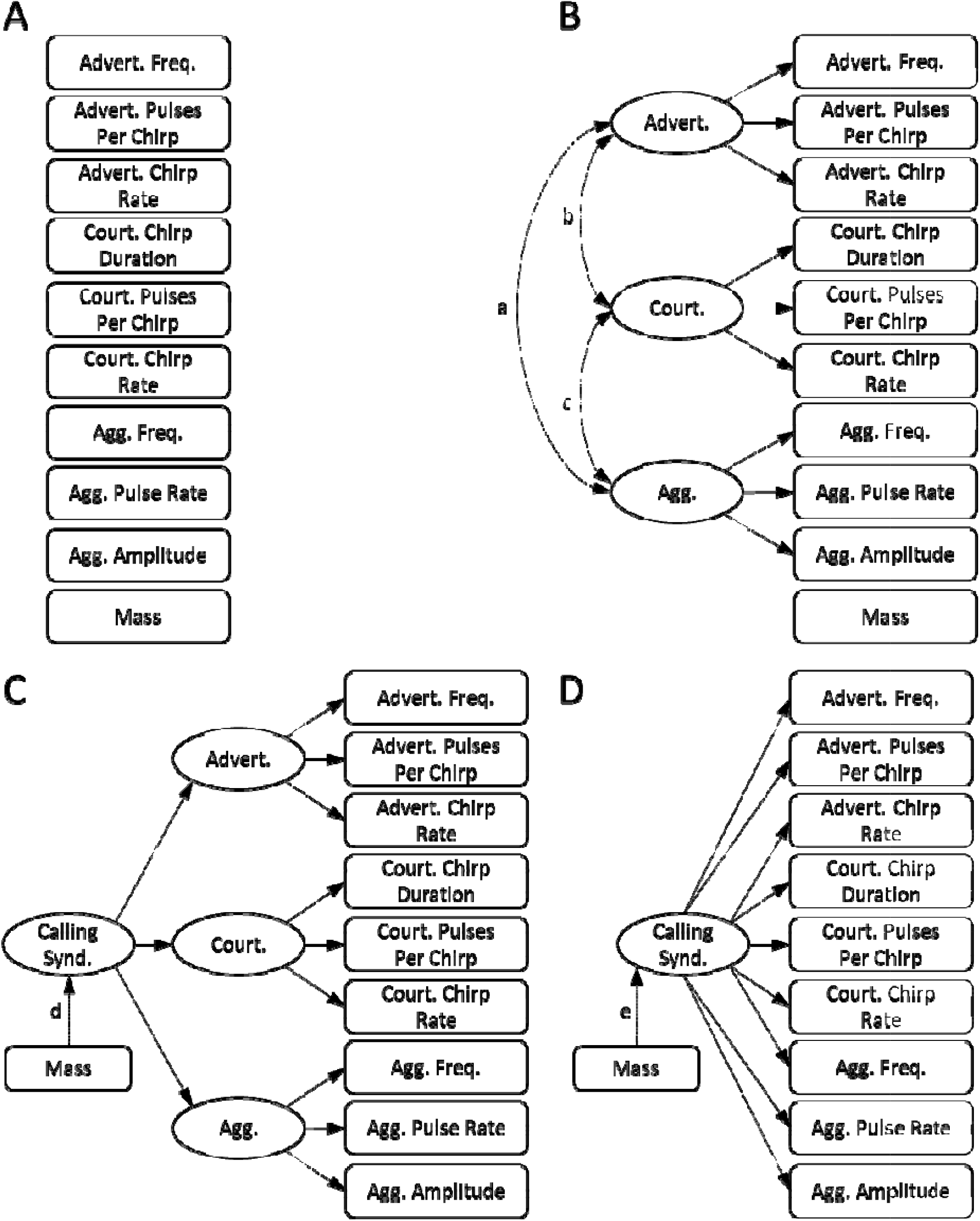
Graphical representations of Structural Equation Models compared. Models are described in the text. Model comparison results are given in Table 3. Single arrows represent causal relationships between a latent variable and call components. Bidirectional arrows represent an undefined correlation between call components.

The among-individual correlation matrices were fit to the following nine *a priori* models (Figure 2):

- Model 1: a null hypothesis where all call components are uncorrelated. Because model comparison approaches are based strictly on those models included, the relative ranking of this model gives an indication of the appropriateness of the suite of models (Figure 2A).
- Models 2-3: two models falling under the classic “Redundant Signal Hypothesis” wherein signals are providing similar information to the listener. In model 2, advertisement and courtship signals are providing females with redundant information about size and quality and advertisement and aggression calls are providing males with redundant information about size and quality (Figure 2B, paths a and b are active—i.e. allowed to be non-zero). In model 3, courtship and aggression calls are also providing redundant information to the receiver (Bertram and Rook 2012) (Figure 2B, paths a, b and c are active). Both of these models correspond to the Functional Modularity model of the systems approach wherein individual signal components serve a similar function but there are overarching functions for each call type.
- Model 4: a model corresponding to a Multiple-Messages Hypothesis wherein all three call types are providing different but complementary information to listening conspecifics (Moller and Pomiankowski 1993, Harrison et al. 2013). This hypothesis corresponds to the Functional Modularity model of the systems approach. The call components of all three call types are uncorrelated in this model (Figure 2B, no lettered paths active).
- Model 5: both courtship and aggression call components will be correlated due to the short range that each call travels (Figure 2B, path c is active). This hypothesis also corresponds to the Functional Modularity model of the systems approach. Courtship and aggression calls are functionally used for short range calling.
- Model 6: Honest Signaling, all of the calling components stem from a single underlying calling phenotype, with no modularity within the three call types. This underlying calling structure is causally affected by mass (Figure 2D, path e is active). This hypothesis corresponds to the Degeneracy model of the systems approach wherein all signal components serve a similar function.
- Model 7: All of the call components stem from a single underlying calling phenotype, with no modularity within the three call types (Figure 2D, no lettered path active). This hypothesis also corresponds to the Degeneracy model under the systems approach terminology.
- Model 8: Honest Signaling with Redundant Signals and Modularity, wherein all three call types causally stem from an underlying calling syndrome or phenotype, implying some signal redundancy. However, each call type still exhibits modularity (i.e. components of aggression calls are more closely related to other components of aggression calls than to components of advertisement calls (Wagner and Hoback 1999, Holzer et al. 2003). This underlying calling structure is causally affected by mass (Figure 2C, path d is active) and so downstream call components are honest signals of mass. This hypothesis corresponds to both the Degeneracy and Functional Modularity models of the systems framework. All signal components serve a similar function, while overarching call types/modules serve different functions, and all of this is causally affected by mass.
- Model 9: Redundant Signals with Modularity, another version of the redundant signals hypothesis wherein all three call types causally stem from an underlying calling syndrome or phenotype, implying some signal redundancy. However, each call type still exhibits modularity. In this model, mass does not affect calling structure (Figure 2C, no lettered path active), i.e. calls do not honestly signal mass. Like model 8, this structure corresponds to both the Degeneracy and Functional Modularity systems definitions.

These models were fit using the `lavaan` package in R and the ability of each model to explain the pattern of correlations compared based on differences of AIC values among models (ΔAIC) (Dingemanse et al., 2010; Dochtermann & Jenkins, 2007). In addition to the nine models listed above, we also tested a “component model” wherein the same components from different call types stem from an underlying component phenotype with no modularity (i.e. courtship pulses per chirp and advertisement pulses per chirp are providing shared information, independent of any specific call types). This model was unfittable, indicating that the model departed too far from the data, suggesting lack of biological relevance.

Because we estimated among-individual correlations using a Markov Chain Monte Carlo approach, we had 1000 estimates of the correlation matrix. Following Araya-Ajoy & Dingemanse (2014) and Dingemanse et al. (2020) the SEM models were fit to each of the 1000 estimated correlation matrices and so we also had 1000 estimates of the AIC and ΔAIC values of each model. Therefore, the model with a ΔAIC whose posterior mode was closest to zero was ranked as the overall best model. This approach allows the estimation of uncertainty around these ΔAIC values so we also considered how often a particular model was ranked as best (i.e. ΔAIC = 0) or could not be distinguished from the best model (i.e. ΔAIC ≤ 2; Dingemanse et al. 2019). We also assessed the “significance” of any particular correlation based on whether its 95% credibility interval overlapped zero (Table S3).

## Results

### Repeatability by Call Type

Advertisement call components were moderately to highly repeatable with peak frequency having the highest adjusted repeatability (τ = 0.58) (Table 2). Components of aggression calls were similarly repeatable, with peak frequency once again having the highest repeatability (τ = 0.48) (Table 2). In contrast to advertisement and aggression call components, courtship call components were generally low to moderately repeatable, with only pulses per chirp (τ = 0.30), chirp duration (τ = 0.30) and chirp rate (τ = 0.38), exhibiting repeatabilites above 0.1 (Table 2). Based on the repeatabilities of all call components estimated from univariate mixed models (Table S3), we used the following components in subsequent analyses: Advertisement – frequency, pulses per chirp and chirp rate, Aggression – frequency, pulse rate, amplitude, Courtship – pulses per chirp, chirp duration and chirp rate (Table 3). Neither the recording chamber in which an individual was recorded nor the box in which an individual was reared contributed substantial variation to any of the call components measured (Table S3). Female ID from courtship trials also did not explain a substantive proportion of the variation present in courtship call components, never explaining more than seven percent of the variation present (Table S3). In contrast, the rival male present explained 5 to 18 % of the variation present in aggression call components (Table S3).

**Table 3.** Model rank along with posterior modal estimates for AIC and ΔAIC values for each model and the number (out of 1000) of MCMC posterior samples for which a particular model was the best fitting (ΔAIC = 0) and or within ΔAIC ≤ 2 from the best fitting model.

| Model number | Model Rank | Classic Signaling Hypothesis | Systems Approach Model | AIC | ΔAIC | AIC = 0 | AIC $\leq 2$ |
| --- | --- | --- | --- | --- | --- | --- | --- |
| 1 | 9 | Null Hypothesis | Null Hypothesis | 2848 | 148 | 0 | 0 |
| 2 | 5 | Redundant Signals | Functional Modularity | 2725 | 25 | 50 | 84 |
| 3 | 3 | Redundant Signals | Functional Modularity | 2718 | 18 | 129 | 224 |
| 4 | 6 | Multiple Messages | Functional Modularity | 2733 | 33 | 8 | 15 |
| 5 | 4 | Short Range Calling | Functional Modularity | 2721 | 21 | 62 | 107 |
| 6 | 8 | Honest Signaling | Degeneracy | 2735 | 35 | 9 | 29 |
| 7 | 7 | Full Integration | Degeneracy | 2734 | 34 | 11 | 30 |
| 8 | 1 | **Honest Signaling and Redundant Signals with Modularity** | **Degeneracy and Functional Modularity** | **2704** | **3** | **545** | **775** |
| 9 | 2 | Redundant Signals with Modularity | Degeneracy and Functional Modularity | 2709 | 9 | 186 | 363 |

### Signal Integration

The best fitting structural equation model corresponded to a model where call components were part of call specific modules (i.e. advertisement vs. aggression vs. courtship calls) and these modules were themselves integrated under a common latent calling syndrome (Model 8; Table 3). This model supports the classic redundant signaling hypothesis as well as both signal degeneracy and functional modularity under the systems approach. Model 9 was also partially supported, with a fit within 2 AIC values over a third of the time (Table 3, Figure 3). These models differed based on whether body mass had a causal effect on call structure, as expected according to the honest signaling hypothesis—with the best fit model supporting this causal link with mass (Table 3, Figures 2 & 3).

**Figure 3.**
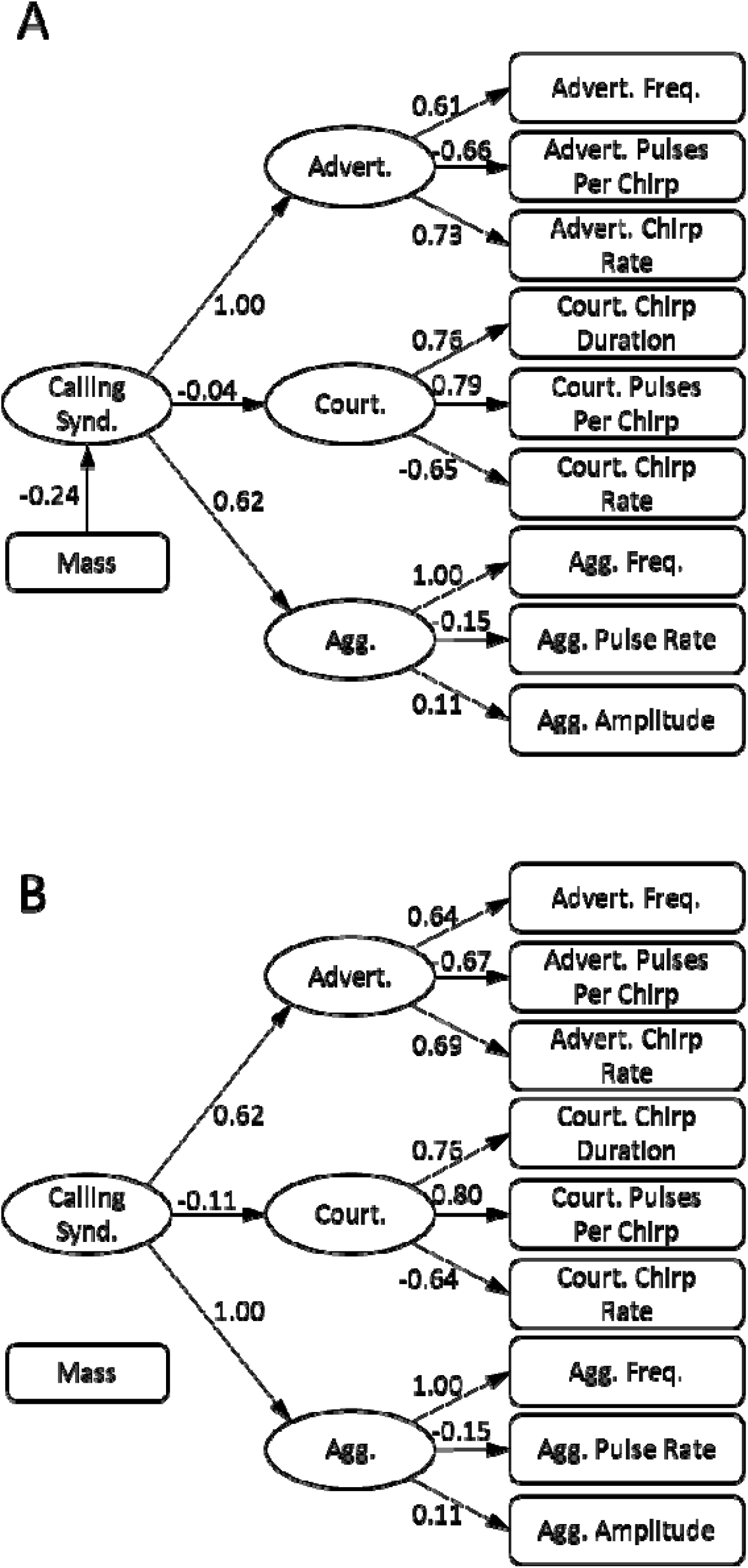
Structural Equation Model Parameter Estimates of the Degeneracy 3 and 4 Models

Overall, the strength of the relationships between the overarching syndrome structure to the call specific latent variables underpinning call components were similar in each model (Figure 3A versus 3B). In both model 8 and 9, all components of advertisement and courtship call were strongly tied to their respective latent variables. In contrast, the aggression latent variable was primarily indicative of call frequency, with limited connections to pulse rate and call amplitude (Figure 3). In addition, the path coefficients connecting the overall calling syndrome latent variable to courtship call approached zero and were substantially weaker than for advertisement and aggression calls (Figures 3A and 3B). This suggests that courtship calls provide independent information not captured by either advertisement or aggression calls.

With the general structure of Model 8 most well supported (Table 3), it is also noteworthy that the relationship between mass and the underlying “Calling Syndrome” was negative (−0.24, Figure 3A). As a result, mass had a negative effect on the expression of the following call components: advert frequency, advert chirp rate, courtship chirp rate, aggression frequency and aggression pulse rate (direction and magnitudes of relationships can be determined by multiplication of path coefficients) (Figure 3A).

## Discussion

Following a systems approach, the two best fit models for *Acheta domesticus* signaling fall under systems definitions of degeneracy and functional modularity. The overall signal syndrome allows different signals to provide similar information (degeneracy) while signal components tie together in lower level modules according to function (functional modularity, Hebets et al. 2016). This result can be further interpreted if the degeneracy hypothesis is integrated with classic signaling hypotheses. Specifically, the Hebets et al. (2016) model of degeneracy closely fits the classic signaling hypotheses of redundant signals, i.e. different signal types provide similar information to receivers (Bertram & Rook, 2012; Harrison et al., 2013). The two best fit models are therefore indicative of an overarching calling syndrome from which the three individual call types stem (see also (Wagner and Hoback 1999, Holzer et al. 2003, Bertram and Rook 2012). The strong link between advertisement and aggression within the overall calling syndrome in the two best fit models (Figure 3) further demonstrates that advertisement and aggression provide similar information about male size and quality (see also Bertram & Rook, 2012). Despite this, the three call types did differ in the strength of association with the calling syndrome and thus level of integration (courtship being the least integrated).

While cricket mass was negatively associated with the calling syndrome, the relationship between mass and specific call components is consistent with previous research on female choice and known morphological relationships with call components. For example, mass was positively related with advertisement pulses per chirp (Figure 3), and higher pulses per chirp preferred by female crickets in other species (Gray 1997). Mass was also negatively related to advertisement and aggression frequency (Figure 3). A negative relationship between the two was previously described result for other cricket species (Simmons and Zuk 1992, Simmons 1995) in which females prefer lower frequencies (Simmons and Ritchie 1996). Consequently, call components likely serve as an honest signal of male body-size in *Acheta domesticus*.

Many of the call components exhibited repeatabilities greater than the average reported for other behaviors (⍰ =0.37, (Bell et al. 2009)). In particular, variation in advertisement call components was strongly explained by individual identity (Tables 1 and S3). These high repeatabilities suggest an upper limit for the heritability of call components in *Acheta domesticus* that would be consistent with heritability of calling in other cricket species (Mousseau and Howard 1998). We used mean and variance standardized values for all call components given those were measured on scales varying on numerous orders of magnitude of difference. While this procedure allowed for proper convergence of our statistical models, one drawback is that additional metrics based on calculating coefficients of variation (Wilson 2008, Dochtermann and Royauté 2019) and evolutionary metrics like trait autonomy are not estimable (Houle 1992, Hansen et al. 2011). How much these integrated patterns of call components are likely to constrain evolutionary responses therefore remains an open question.

In further examining other contributors to variation in call components, one particularly interesting finding is that variation in courtship call components was not substantially influenced by female receiver identity (Table S3). In other words, the identity of the female being courted did not affect male behavior. This indirectly suggests that males do not modify their own courtship based on differences amongst females, an interesting topic for future research. In contrast to female identity, rival male identity substantially influenced the variation in both amplitude and frequency of aggression calls (Table S3). This makes intuitive sense: a higher quality opponent may deter a lower quality male from producing a strong call. These findings also align with previous research of indirect genetic effects on aggression in other Gryllid species (Santostefano et al. 2016, Santostefano et al. 2017). Finally, despite having accounted for multiple variables involved in courtship expression, more than 55 % of the phenotypic variation stems from unexplained sources of within-individual variation (Table S3). This within-individual variation corresponds to variation arising from temporary environmental effects and are in line with the findings of Harrison et al. (2013) showing a curvilinear relationship between residual mass and courtship pulse and chirp rates wherein both small and large crickets used similar pulse and chirp rates to court females (Harrison et al. 2013). These results suggests that we still know little about what courtship is communicating to females. Further exploration of environmental contributions to courtship plasticity is therefore needed regarding signaling hypotheses for the courtship calls of crickets.

Studies evaluating integration and modularity among sexual signaling traits are increasingly common with continued and increasing interest in multimodal communication. For example, in the black field cricket (*Teleogryllus commodus*), call components were found to be highly integrated and vary little among populations or diet quality (Pitchers et al. 2013). Song components in barn swallows have also been found to be tightly integrated but relatively independent from morphology and coloration modules (Wilkins et al. 2015). In contrast, the degree of integration between advertising and aggressive calls varies substantially among species of treefrogs (Reichert and Hobel 2018). The pattern of integration found here (Figure 3) suggests support for both honest signaling as well as degeneracy, depending on which approach is implemented, where particular components of a call type are more strongly related to each other than across call types. Phenotypic integration can occur in a strictly modular fashion or integration can exist with modularity, due to shared developmental pathways or functions (Araya-Ajoy and Dingemanse 2014, Royaute et al. 2015). Such integration with underlying modularity, like that described here, has been termed “quasi-independent modularity” (Larouche et al. 2018). Quasi-independent modularity results in an inability of constituent components to evolve independently but, counter-intuitively, can also facilitate rapid evolutionary change (West-Eberhard 2003).

Unfortunately, much of the literature on signaling has focused on phenotypic variation which conflates multiple sources of variation (Adolph and Hardin 2007, Dingemanse et al. 2012). Our attempt here to evaluate correlations of call components at the level of among-individual (co)variance should more closely capture the underlying genetic structure of call integration (Dingemanse and Dochtermann 2014, Dochtermann et al. 2015). To the degree that among-individual covariation captures genetic covariation, our results are consistent with advertisement and aggression calls being genetically integrated while courtship calls are their own quasi-independent genetic module. This suggests that advertisement and aggression calls will likely evolve in a correlated manner while courtship calls have the capacity to follow a more independent evolutionary trajectory.

The quasi-independent structure that we identified for *Acheta domesticus* combines two categories of the systems framework proposed by Hebets et al. (2016): degeneracy and functional modularity (Figure 2, Table 3). This observed structure is also consistent with hypotheses of redundant signaling and honest signaling. While the systems approach provides more precise mechanistic descriptions of signaling, the classical hypotheses of signaling are more explicitly linked to evolutionary mechanisms and sexual selection.

Having two competing conceptual frameworks, particularly when they have overlapping terminology, has the potential to hinder our understanding of communication. Fortunately, these two frameworks can be combined to offer greater insight (Table 1). For example here, cricket calling’s degenerate and functional modularity is best described by the systems approach and the connection to mass is best described by classic hypotheses of honest signaling. Combining these two frameworks suggests that cricket calls are structured according to a pattern of honest integration (Table 1). By highlighting this important disconnect between both the classic and modern signaling frameworks as well as similarities between the two frameworks, we have shown that the two frameworks can be integrated together. This combination contributed here to a better understanding of pre-mating signaling phenotypes and of patterns of signal integration.

## Acknowledgements

We thank M. Anderson Berdal, J. Dalos, J. Albers, K. Pnewski, M. Rick, T. Tesarek and A. Wilson for assistance with cricket rearing and data collection. We also thank E. Gillam for assistance with cricket calling analysis. This study was funded by NSF IOS-1557951 (to NAD) and the Department of Biological Sciences at North Dakota State University.

## Supporting Information for

**Table S1.**
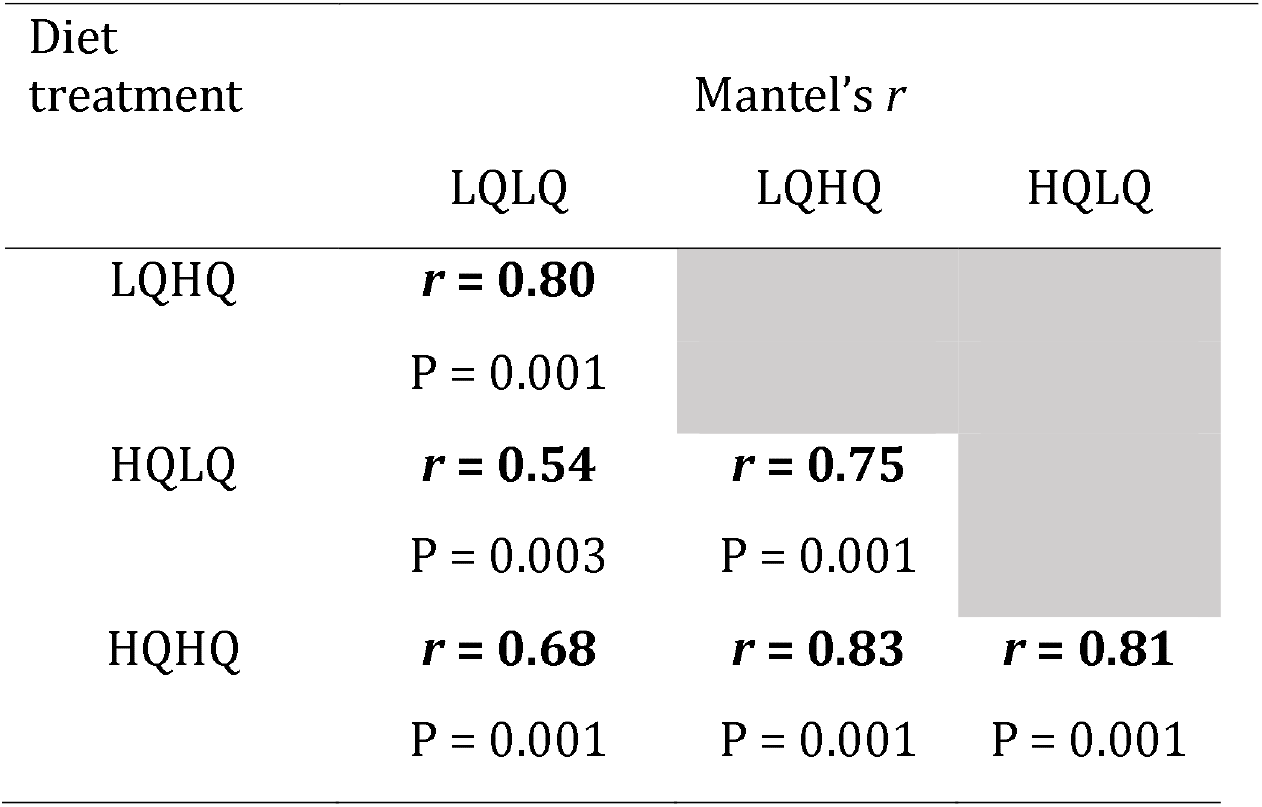
Mantel’s tests for comparing correlation matrices similarity across diet treatments. The Mantel’s *r* represents how closely related the among correlation matrices are between two diets. Bold indicates statistically significant correlations.

**Table S2.** Number of individuals who went through each repetition of recording for each call type

| **Repetition** | **Advertisement** | **Aggression** | **Courtship** | **Total** |
| --- | --- | --- | --- | --- |
| **1** | 127 | 91 | 105 | 323 |
| **2** | 122 | 87 | 101 | 310 |
| **3** | 107 | 85 | 100 | 292 |
| **4** | 5 |  |  | 5 |

**Table S3.**
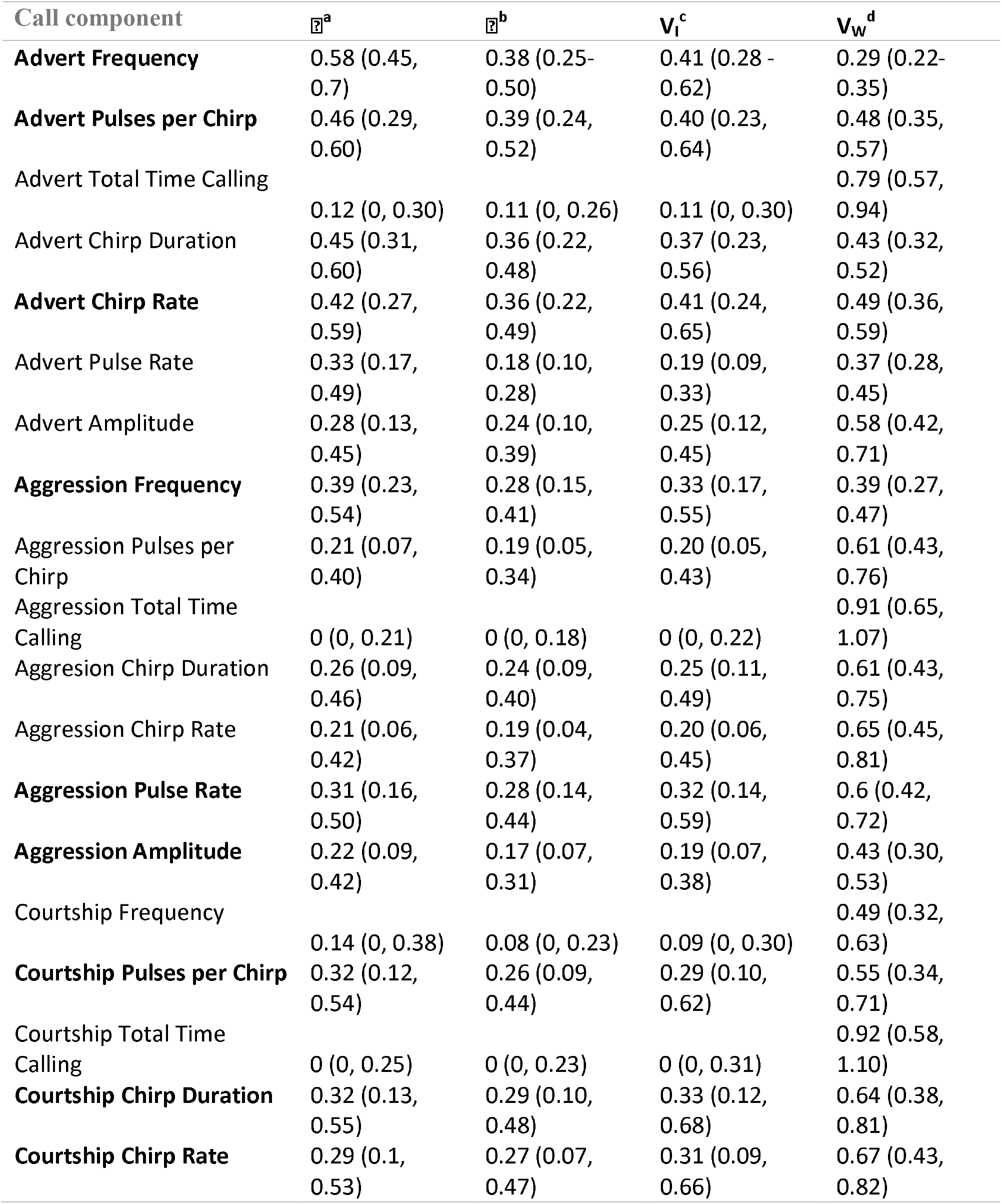
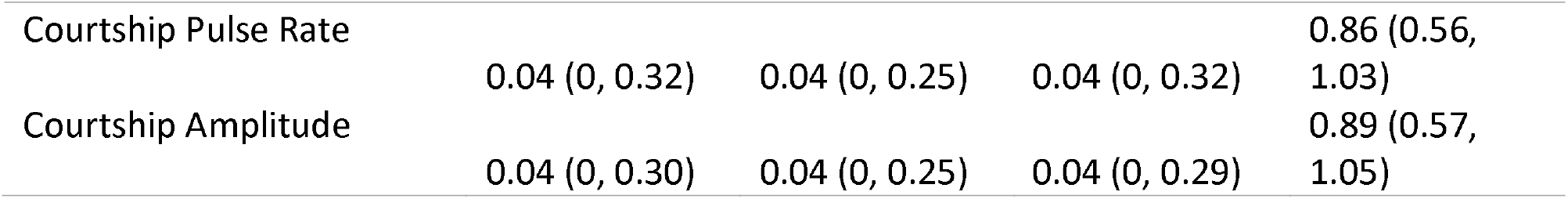
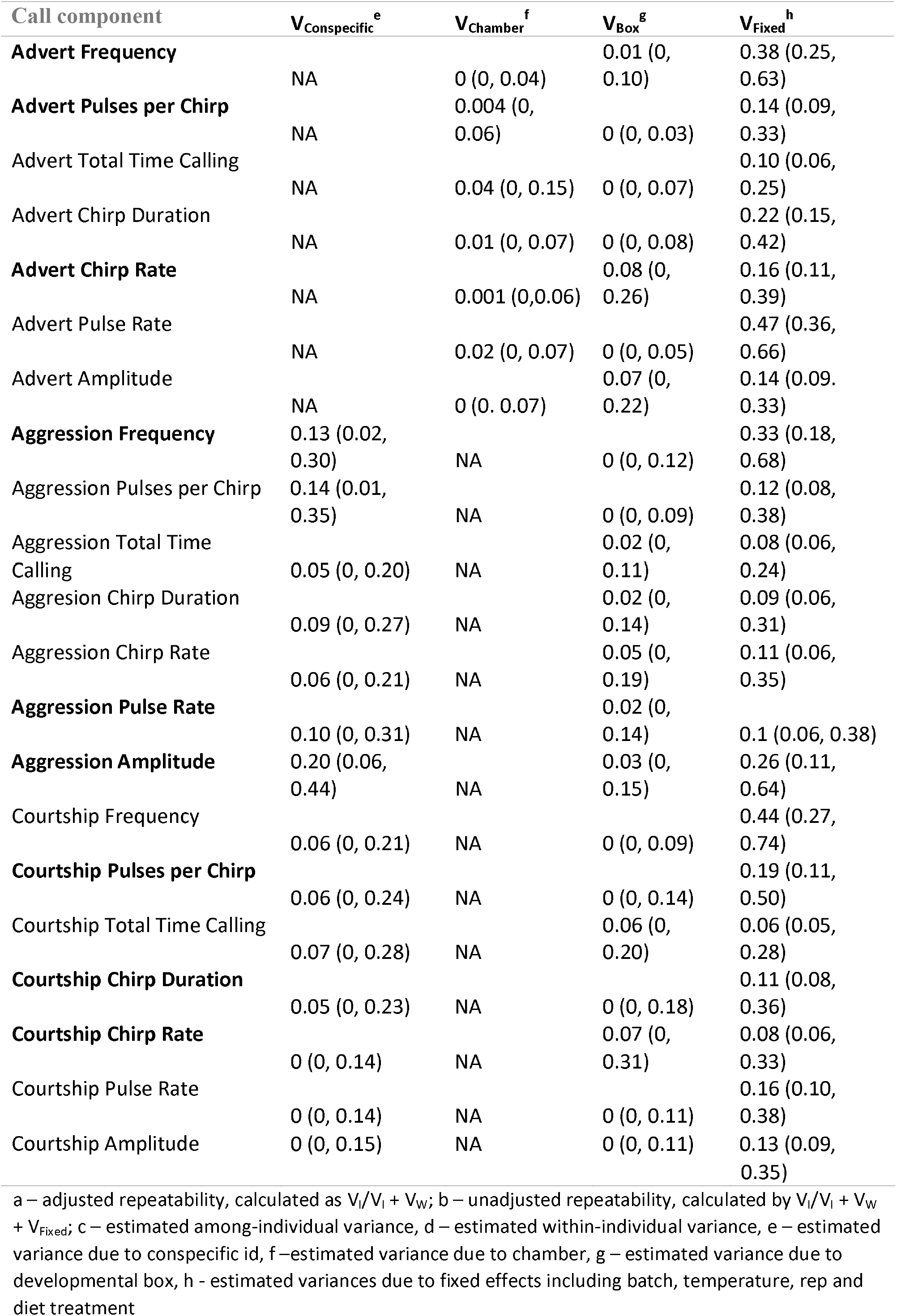
Repeatabilities and variances at the among- and within-individual levels as well as the fixed and random effects for each call parameter calculated using the rptR package with 95% confidence intervals. The bolded call components represent those chosen for the condensed multi-response MCMCglmm model

**Table S4.** Among and within-individual correlations, along with repeatabilities, as estimated by a multi-response mixed effects model. Among-individual correlations are above the diagonal, within-individual correlations are below the diagonal, with repeatabilities (calculated from the multi-response model rather than the univariate models of Table 2) shown on the diagonal. Shaded within-individual correlations are inestimable and overlap with zero. 95% credibility intervals for repeatabilities are reported along the diagonal. Bold values are the among- and within-individual correlations whose 95% credibility intervals did not overlap zero.

|  | Advert_<br>FQ | Advert_<br>PC | Advert_<br>CR | Court_C<br>D | Court_<br>PC | Court_C<br>R | Agg_F<br>Q | Agg_P<br>R | Agg_A<br>mp | Mass |
| --- | --- | --- | --- | --- | --- | --- | --- | --- | --- | --- |
| Advert_<br>FQ | 0.64<br>(0.53-0.73) | -0.09 | 0.09 | -0.09 | -0.13 | 0.14 | **0.41** | 0.00 | -0.12 | -<br>0.18 |
| Advert_<br>PC | 0.12 | 0.50<br>(0.39-0.62) | **-0.58** | 0.24 | 0.16 | -0.23 | -0.08 | -0.06 | 0.21 | 0.09 |
| Advert_<br>CR | **-0.17** | **-0.71** | 0.58<br>(0.42-0.73) | -0.20 | -0.22 | **0.30** | 0.21 | 0.25 | -0.28 | -<br>0.21 |
| Court_C<br>D |  |  |  | 0.47<br>(0.32-0.60) | **0.62** | **-0.48** | -0.03 | **-0.33** | 0.21 | 0.15 |
| Court_P<br>C |  |  |  | **0.81** | 0.48<br>(0.34-0.61) | **-0.41** | 0.00 | -0.27 | 0.22 | 0.08 |
| Court_C<br>R |  |  |  | **-0.54** | **-0.49** | 0.48<br>(0.34-0.63) | 0.10 | 0.18 | -0.14 | -<br>0.19 |
| Agg_FQ |  |  |  |  |  |  | 0.53<br>(0.39-0.63) | -0.09 | 0.05 | -<br>0.13 |
| Agg_PR |  |  |  |  |  |  | **-0.24** | 0.45<br>(0.35-0.62) | -0.29 | -<br>0.11 |
| Agg_Am<br>p |  |  |  |  |  |  | **0.42** | **-0.27** | 0.45<br>(0.32-0.55) | 0.20 |
| Mass |  |  |  |  |  |  |  |  |  | 0.48<br>(0.3<br>5-0.68<br>) |

FQ - frequency, PC – pulses per chirp, CR – chirp rate, CD – chirp duration, PR – pulse rate, Amp - amplitude

## References

Adolph, S. C., and J. S. Hardin. 2007. Estimating phenotypic correlations: correcting for bias due to intraindividual variability. Functional Ecology 21:178–184.

Andersson, M., and L. W. Simmons. 2006. Sexual selection and mate choice. Trends in Ecology & Evolution.

Araya-Ajoy, Y. G., and N. J. Dingemanse. 2014. Characterizing behavioural ‘characters’: an evolutionary framework. Proceedings of the Royal Society B-Biological Sciences 281.

Bell, A. M., S. J. Hankison, and K. L. Laskowski. 2009. The repeatability of behaviour: a meta-analysis. Animal Behaviour 77:771–783.

Berglund, A., A. Bisazza, and A. Pilastro. 1996. Armaments and ornaments: An evolutionary explanation of traits of dual utility. Pages 385–399, Biological Journal of the Linnean Society.

Bertram, S. M., and V. Rook. 2012. Relationship between condition, aggression, signaling, courtship, and egg laying in the field cricket, Gryllus assimilis. Ethology 118:360–372.

Bro-Jørgensen, J., T. Dabelsteen, and J. a. Email authorBro-Jørgensen. 2008. Knee-clicks and visual traits indicate fighting ability in eland antelopes: multiple messages and back-up signals. BMC Biology 6.

Buchanan, K. L., K. A. Spencer, A. R. Goldsmith, and C. K. Catchpole. 2003. Song as an honest signal of past developmental stress in the European starling (Sturnus vulgaris). Proceedings of the Royal Society B-Biological Sciences 270:1149–1156.

Dingemanse, N. J., I. Barber, and N. A. Dochtermann. 2019. Non-consumptive effects of predation: does perceived risk strengthen the genetic integration of behaviour and morphology in stickleback? Ecology Letters.

Dingemanse, N. J., and N. A. Dochtermann. 2014. Individual behaviour: behavioural ecology meets quantitative genetics. Quantitative Genetics in the Wild:54–67.

Dingemanse, N. J., N. A. Dochtermann, and S. Nakagawa. 2012. Defining behavioural syndromes and the role of ‘syndrome deviation’ in understanding their evolution. Behavioral Ecology and Sociobiology 66:1543–1548.

Dingemanse, N. J., N. A. Dochtermann, and J. Wright 2010. A method for exploring the structure of behavioural syndromes to allow formal comparison within and between data sets. Animal Behaviour 79:439–450.

Dochtermann, N. A., and S. H. Jenkins. 2007. Behavioural syndromes in Merriam’s kangaroo rats (*Dipodomys merriami*): a test of competing hypotheses. Proceedings Of The Royal Society B-Biological Sciences 274:2343–2349.

Dochtermann, N. A., and R. Royauté. 2019. The mean matters: going beyond repeatability to interpret behavioural variation. Animal Behaviour 153:147–150.

Dochtermann, N. A., T. Schwab, and A. Sih. 2015. The contribution of additive genetic variation to personality variation: heritability of personality. Proceedings of the Royal Society B-Biological Sciences 282.

Doucet, S. M., and R. Montgomerie. 2003. Multiple sexual ornaments in satin bowerbirds: ultraviolet plumage and bowers signal different aspects of male quality Behavioral Ecology 14:503–509.

Falconer, D. S. 1996. Introduction to Quantitative Genetics. Pages 122–145 *in* T. F. C. Mackay, editor. Pearson Education Limited.

Garrison, C. 2017. Environmental Components of Phenotypic Variation: Dietary and Trans-generational Effects on Behavior. North Dakota State University, ProQuest Dissertations and Theses.

Gray, D. A. 1997. Female house crickets, Acheta domesticus, prefer the chirps of large males. Animal Behaviour 54:1553–1562.

Gray, D. A., and G. Eckhardt. 2001. Is cricket courtship song condition dependent? Animal Behaviour 62:871–877.

Hadfield, J. D. 2010. MCMC Methods for Multi-Response Generalized Linear Mixed Models: The MCMCglmm R Package. Journal of Statistical Software 33:1–22.

Hansen, T. F., C. Pelabon, and D. Houle. 2011. Heritability is not evolvability. Evolutionary Biology 38:258.

Harrison, S. J., I. R. Thomson, C. M. Grant, and S. M. Bertram. 2013. Calling, Courtship, and Condition in the Fall Field Cricket, Gryllus pennsylvanicus. Plos One 8:9.

Hebets, E., and D. Papaj. 2005. Complex signal function: developing a framework of testable hypotheses. Behavioral Ecology and Sociobiology 57:197–214.

Hebets, E. A., A. B. Barron, C. N. Balakrishnan, M. E. Hauber, P. H. Mason, and K. L. Hoke. 2016. A systems approach to animal communication. Proceedings of the Royal Society B-Biological Sciences.

Hedrick, A., and T. Weber. 1998. Variance in female responses to the fine structure of male song in the field cricket, Gryllus integer. Behavioral Ecology 9:582–591.

Hedrick, A. V. 1986. Female preferences for male calling bout duration in a field cricket. Behavioral Ecology and Sociobiology 19:73–77.

Hedrick, A. V. 1988. Female choice and the heritability of attractive male traits: an empirical study. The American Naturalist 132:267–276.

Holzer, B., A. Jacot, and M. W. G. Brinkhof. 2003. Condition-dependent signaling affects male sexual attractiveness in field crickets, Gryllus campestris. Behavioral Ecology 14:353–359.

Houle, D. 1992. Comparing evolvability and variability of quantitative traits. Genetics 130:195–204.

Johnstone, R. A. 1996. Multiple displays in animal communication: ‘Backup signals’ and ‘multiple messages’. Philosophical Transactions of the Royal Society of London Series B-Biological Sciences 351:329–338.

Larouche, O., M. L. Zelditch, and R. Cloutier. 2018. Modularity promotes morphological divergence in ray-finned fishes. Scientific Reports 8.

Moller, A. P., and A. Pomiankowski. 1993. WHY HAVE BIRDS GOT MULTIPLE SEXUAL ORNAMENTS. Behavioral Ecology and Sociobiology 32:167–176.

Moreno-Gomez, F. N., L. D. Bacigalupe, A. A. Silva-Escobar, and M. Soto-Gamboa. 2015. Female and male phonotactic responses and the potential effect of sexual selection on the advertisement calls of a frog. Pages 79–86, Animal Behavior.

Mousseau, T., and D. Howard. 1998. Genetic variation in cricket calling song across a hybrid zone between two sibling species. Evolution 52:1104–1110.

Nakagawa, S., and H. Schielzeth. 2010. Repeatability for Gaussian and non-Gaussian data: a practical guide for biologists. Biological reviews of the Cambridge Philosophical Society 85:935–956.

Pigliucci, M. 2003. Phenotypic integration: studying the ecology and evolution of complex phenotypes. Ecology Letters 6:265–272.

Pitchers, W. R., R. Brooks, M. D. Jennions, T. Tregenza, I. Dworkin, and J. Hunt. 2013. Limited plasticity in the phenotypic variance-covariance matrix for male advertisement calls in the black field cricket, Teleogryllus commodus. Journal of Evolutionary Biology 26:1060–1078.

Reichert, M. S., and G. Hobel. 2018. Phenotypic integration and the evolution of signal repertoires: A case study of treefrog acoustic communication. Ecology and Evolution 8:3410–3429.

Rosenthal, M. F., M. R. Wilkins, D. Shizuka, and E. A. Hebets. 2018. Dynamic changes in display architecture and function across environments revealed by a systems approach to animal communication. Pages 1134–1145, Evolution.

Royauté, R., C. Garrison, J. Dalos, M. A. Berdal, and N. A. Dochtermann. 2019. Current energy state interacts with the developmental environment to influence behavioural plasticity. Animal Behaviour 148:39–51.

Royauté, R., and N. A. Dochtermann. 2017. When the mean no longer matters: developmental diet affects behavioral variation but not population averages in the house cricket (*Acheta domesticus*). Behavioral Ecology 28:337–345.

Royaute, R., K. Greenlee, M. Baldwin, and N. A. Dochtermann. 2015. Behaviour, metabolism and size: phenotypic modularity or integration in Acheta domesticus? Animal Behaviour 110:163–169.

Santostefano, F., A. J. Wilson, Y. G. Araya-Ajoy, and N. J. Dingemanse. 2016. Interacting with the enemy: indirect effects of personality on conspecific aggression in crickets. Behavioral Ecology 27:1235–1246.

Santostefano, F., A. J. Wilson, P. T. Niemela, and N. J. Dingemanse. 2017. Indirect genetic effects: a key component of the genetic architecture of behaviour. Scientific Reports 7.

Schluter, D., and T. Price. 1993. HONESTY, PERCEPTION AND POPULATION DIVERGENCE IN SEXUALLY SELECTED TRAITS. Proceedings of the Royal Society B-Biological Sciences 253:117–122.

Simmons, L. W. 1995. Correlates of male quality in the field cricket, *Gryllus campestris* L.: age, size, and symmetry determine pairing success in field populations. Behavioral Ecology 6:376–381.

Simmons, L. W., and M. G. Ritchie. 1996. Symmetry in the songs of crickets. Proceedings of the Royal Society of London, Series B 263:1305–1311.

Simmons, L. W., and M. Zuk. 1992. Variability in call structure and pairing success of male field crickets, *Gryllus bimaculatus:* the effects of age, size and parasite load. Amnimal Behaviour 44:1145–1152.

Spencer, K. A., K. L. Buchanan, A. R. Goldsmith, and C. K. Catchpole. 2003. Song as an honest signal of developmental stress in the zebra finch (Taeniopygia guttata). Hormones and Behavior 44:132–139.

Stoffel, M., S. Nakagawa, and H. Schielzeth. 2017. rptR: Repeatability estimation and variance decomposition by generalized linear mixed-effects models. Methods in Ecology and Evolution.

Wagner, W. E., and W. W. Hoback. 1999. Nutritional effects on male calling behaviour in the variable field cricket. Animal Behaviour 57:89–95.

West-Eberhard, M. J. 2003. Developmental Plasticity and Evolution. Oxford University Press.

Wilkins, M. R., D. Shizuka, M. B. Joseph, J. K. Hubbard, and R. J. Safran. 2015. Multimodal signalling in the North American barn swallow: a phenotype network approach. Proceedings of the Royal Society B-Biological Sciences 282.

Wilson, A. 2008. Why h2 does not always equal VA/VP? Journal of Evolutionary Biology 21:647–650.

Woodgate, J. L., M. M. Mariette, A. T. D. Bennett, S. C. Griffith, and K. L. Buchanan. 2012. Male song structure predicts reproductive success in a wild zebra finch population. Animal Behaviour 83:773–781.

Zahavi, A. 1975. MATE SELECTION - SELECTION FOR A HANDICAP. Journal of Theoretical Biology 53:205–214.

Zuk, M., D. Rebar, and S. P. Scott. 2008. Courtship song is more variable than calling song in the field cricket Teleogryllus oceanicus. Animal Behaviour 76:1065–1071.

